# Dynamic vocal learning in adult marmoset monkeys

**DOI:** 10.1101/2023.09.22.559020

**Authors:** Nikhil Phaniraj, Kaja Wierucka, Judith M. Burkart

## Abstract

While vocal learning is vital to language acquisition in children, adults continue to adjust their speech while adapting to different social environments in the form of social vocal accommodation (SVA). Even though adult and infant vocal learning seemingly differ in their properties, whether the mechanisms underlying them differ remains unknown. The complex structure of language creates a challenge in quantifying vocal changes during SVA. Consequently, animals with simpler vocal communication systems are powerful tools for understanding the mechanisms underlying SVA. Here, we tracked acoustic changes in the vocalizations of adult common marmoset pairs, a highly vocal primate species known to show SVA, for up to 85 days after pairing with a new partner. We identified four properties of SVA in marmosets: (1) bidirectional learning, (2) exponential decrease in vocal distance with time, (3) sensitivity to initial vocal distance, and (4) dyadic acoustic feature synchrony. We developed a mathematical model that shows all four properties. The model suggests that marmosets continuously update the memory of their partners’ vocalizations and modify their own vocalizations to match them, a dynamic form of vocal learning. The model provides crucial insights into the mechanisms underlying SVA in adult animals and how they might differ from infant vocal learning.

## Introduction

Human language, the most complex vocal communication system among animals, is grounded in vocal production learning (VPL) – the ability to alter one’s vocalization properties in response to hearing other vocalizations^1–3^. The past few decades of VPL research has had a strong focus on songbirds, the first non-human clade in which this trait was discovered^4^. Many juvenile songbirds learn their songs during the early days of their life from a tutor, similar to how human infants learn speech from adults^5^. It has been hypothesized that songbirds can do so by memorizing the tutor’s song, forming an auditory template, and matching their own songs to the template^6,7^. This hypothesis and the various neurobiological experiments that support it (reviewed in^8,9^) provide insights into the ways in which nervous systems can achieve VPL.

Even though the most striking example of VPL with the most drastic changes in vocalizations is speech development in human infants, human adults continue to show VPL as they modify their speech properties to adapt to changing social environments. This is visible as social vocal accommodation (SVA), a form of VPL where human interlocutors converge in speech properties such as fundamental frequency, speech rate, word choice and pronunciation, among others^10–12^. A major feature that distinguishes SVA from VPL shown by human infants and juvenile songbirds is that learning is largely unidirectional in the latter, i.e., infants learn to imitate adult vocalizations. In contrast, learning during SVA can be bidirectional, where both isnterlocutors modify speech properties, leading to vocal convergence. This suggests differences in the nature of the processes underlying VPL in infants and adults.

The processes underlying SVA in humans remain largely unknown as the complex structure of language creates an enormous challenge for revealing the mechanisms. As in the case of speech development, SVA research would highly benefit from an animal model with a relatively simple vocal communication system eliciting a similar phenomenon. Common marmosets (*Callithrix jacchus*) stand out among non-human primates as a compelling model due to their remarkable vocal flexibility, which is based in their highly social nature and a cooperative breeding system^13–16^. Marmosets have been shown to possess dialects^17^, and translocation experiments have confirmed the presence of SVA in different call types^18,19^. In this study, we took advantage of this system to comprehensively analyze the temporal dynamics of SVA and arrive at a mathematical model explaining its mechanisms. To study the temporal dynamics of SVA, we used the marmoset SVA dataset collected by Zürcher et al. ^18^ and chose 7 pairs for which recordings with high temporal resolution were available (n=14 marmosets: 7 males and 7 females). Adult marmoset individuals of the opposite sex, originally from groups possessing different dialects ^17^, were paired. Their vocalizations were recorded before pairing, and repeatedly after pairing, until approximately 60 days after pairing (61 ± 16 days, median ± std.) and overall, all vocalizations (trills, phees and food calls) converged over time (18). However, as noted by Cohen-Priva and Sanker^20^, current methods for quantifying SVA may suffer from underestimation due to poor selection of acoustic features whose baseline levels are already similar in the beginning among interlocutors. To overcome this and provide a highly accurate description of the amount and temporal dynamic of marmoset SVA, we used the top 20 acoustic features that best distinguished the calls of the individuals out of a total of 3255 features, as determined in Phaniraj et al.^21^ using a machine-learning classifier (Table S1). To quantify vocal accommodation, for each pair, we measured the Euclidean distance between the centroids (in the acoustic feature space) of all trill calls obtained from the two individuals before and after pair formation. To study the temporal dynamics of SVA we focused on trills because (1) the extent of SVA was highest in trills^19^, and (2) they are close-distance contact calls and are therefore always directed towards the partner. We analyzed a total of 5842 trills across the 14 individuals (1088 before pairing and 4754 after pairing).

In the first step, we analyzed the directionality of VPL in each dyad and how the vocal distance between the dyads changed over time. Moreover, preliminary analyses on a smaller trill dataset suggested that, over time, dyads show synchronized changes in some call parameters, such as fundamental frequency^22^. We, therefore, also analyzed to what extent changes in call structure are synchronized within a dyad using Fréchet distance^23^ and Dynamic Time Warping measures^24^.

In the second step, we compiled four biologically plausible mathematical models that could explain the patterns of SVA in marmosets. The first model, the *Initial Auditory Template Matching (IATM)* was inspired by Konishi’s model for birdsong learning^25^ and assumes that marmosets would form an initial, static auditory template of the partner’s call and match their own calls to the static template (Fig. 1A). The second model, the *Convergence to Intermediate Value (CIV)* was inspired by the preactive and latent auditory templates hypothesized by Marler and Nelson^26^ and assumes that in addition to the initial auditory template of the partner’s call (static acquired template), marmosets possess a static internal template of their own call, which may have previously played a role in vocal development, and both these templates drive VPL (Fig. 1B). We build the third model, the *Dynamic Auditory Template Matching (DATM)* over *IATM* and assume that the auditory template is dynamic and takes into account any changes in the partner’s vocalizations (Fig. 1C). Similar to *DATM*, the fourth model, the *Dynamic Convergence to Intermediate Value (DCIV)* is built upon a previous model, the *CIV*, and assumes that both the internal and acquired templates are dynamic (Fig. 1D). We found the dynamic models *(DATM and DCIV)* to best explain the properties of marmoset SVA. We finally confirm this by simulating SVA in virtual marmoset pairs using the dynamic model and comparing its properties to the empirical data.

**Fig. 1.**
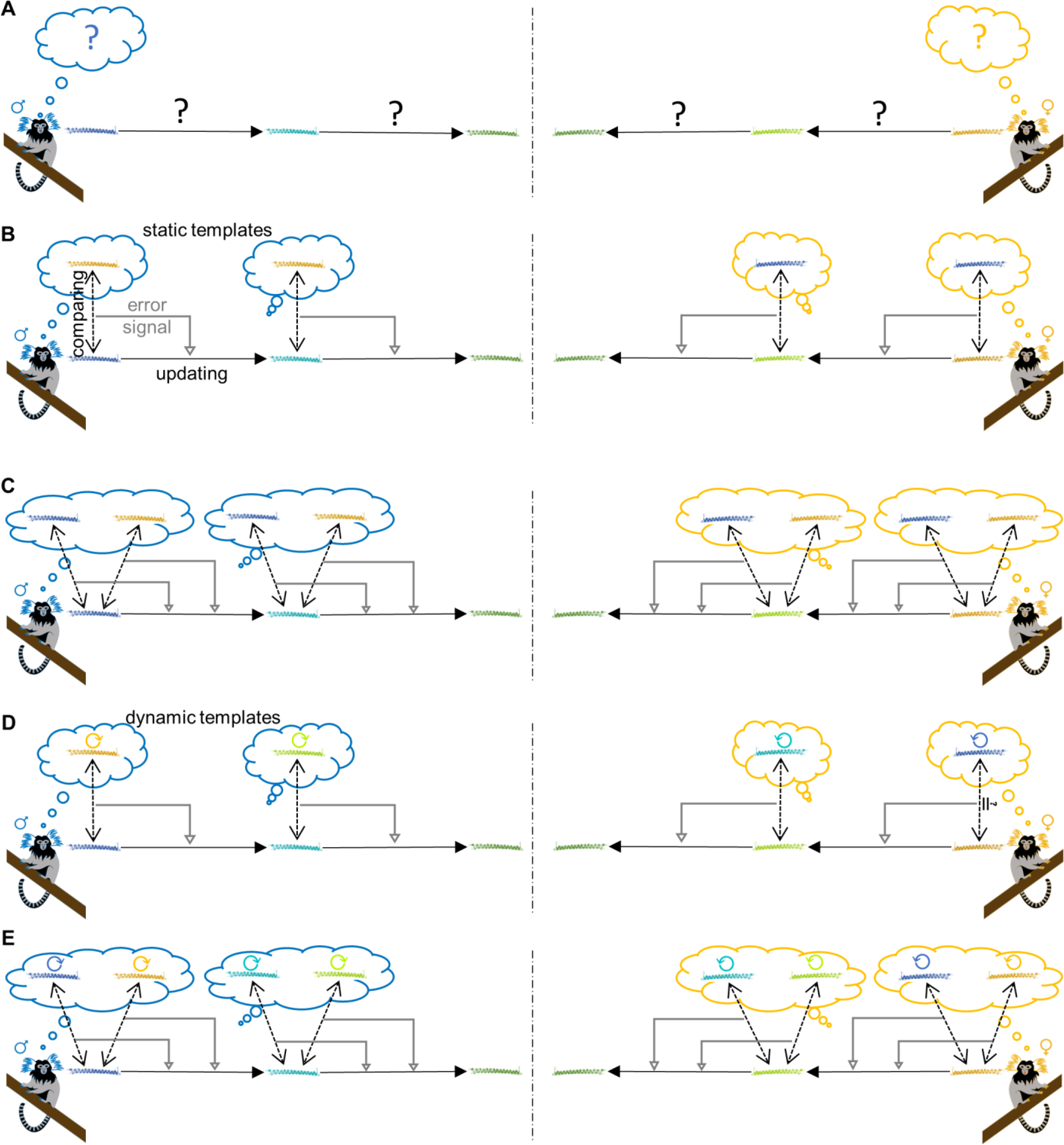
Marmoset vocal accommodation and its models. (**A**) Marmoset vocal accommodation. When marmosets are paired, the trill calls of the individuals become more similar over time. What drives these changes and how is not known. (**B**) Initial Auditory Template Matching (IATM). When paired, marmosets form a static auditory template of the trills of their partner. They compare their own trills to the static template, producing an error signal. This error signal drives VPL as the marmosets modify their trills to reduce the magnitude of the error signal. (**C**) Convergence to Intermediate Value (CIV). When paired, marmosets form a static auditory template of the trills of their partner (static acquired template) in addition to the static internal template that they already possess. Their trills are compared to both the templates and the resulting error signals drive changes in the marmoset’s own trills. (**D**) Dynamic Auditory Template Matching (DATM). When paired, marmosets form an auditory template of the trills of their partner. Marmosets compare their own trills to the template and the error signal drives changes in their own trills. Marmosets then continuously update the template (dynamic template) to account for changes in their partner’s trills, hence producing a dynamic error signal that drives VPL. (**E**) Dynamic Convergence to Intermediate Value (DCIV). Marmosets acquire a template of their partner’s latest trill in addition to the internal template when paired. Both these templates are dynamic, hence producing dynamic error signals when their own calls are compared to both templates, which then drives VPL. In all the panels, black dashed double arrows depict the comparison of the monkey’s current trill with the template(s). Grey arrows represent the error signals. Solid arrows represent the update or change occurring to the trills.

## Results

### Properties of SVA in trills

We noted four characteristics of SVA in marmoset trill calls (Fig. 2A). **(1) Bidirectional learning:** males and females underwent equal amounts of vocal change during SVA (Fig. 2D, n = 7 pairs, t = -0.094, df = 6, p = 0.928, two-tailed paired t-test), hence confirming that learning during marmoset SVA is bidirectional. Therefore, in this respect, VPL in adult marmosets is similar to adult humans and different from human infants and juvenile songbirds. **(2) Exponential decrease of vocal distance with time:** the vocal distance between pairs decayed exponentially after pairing (Fig. 2B). **(3) Sensitivity to initial vocal distance:** the amount of SVA undergone by the pair was significantly positively correlated to the initial vocal distance between them before pairing (Fig. 2C, Spearman’s correlation coefficient r = 0.89, p = 0.01). **(4) Dyadic acoustic feature synchrony:** we developed two ways to measure synchronized changes in acoustic features of calls of dyads over time that were independent of the amount of convergence between individuals using Dynamic Time Warping (DTW) and Fréchet distance measures. Synchrony was found to be significantly higher in actual pairs than in control pairs (i.e. all possible combinations of different-sex dyads in the data set), visible as lower Fréchet and DTW distance values (n=7 actual pairs and 42 control pairs; Fig. 2E, t-value = -4.64, df = 7.06, p = 0.0023 for Fréchet distance; Fig S1A, t-value = -3.02, df = 7.16, p = 0.0188 for DTW distance; two-tailed Welch’s t-test). We visualized this synchrony by reducing the 20-dimensional feature space to 3 principal components that explained most of the variation and plotting the vocal trajectories taken by the male and the female in a pair with time (Movie S1). The DTW and Fréchet distance values of pairs were highly correlated (Fig. S1B, n = 7 pairs, r = 0.94, p = 0.02, Pearson’s correlation). Note that this dyadic acoustic feature synchrony is also apparent when the amount of convergence between individuals is not accounted for (Fig. S2A, n = 7 actual pairs and 42 control pairs, t-value = -8.09, df = 6.42, p = 1.35e-04, two-tailed Welch’s t-test) and while looking at dyadic phase coherence of the principal component of commonly used spectral features for analyzing marmoset vocalizations (mean fundamental frequency, mean spectral entropy, frequency of amplitude modulation and call duration) (Fig. S2B, n = 7 actual pairs and 42 control pairs, t-value = 2.27, df = 9.61, p = 0.047, two-tailed Welch’s t-test) and fundamental frequency of the call alone (Fig. S2C, n = 7 actual pairs and 42 control pairs, t-value = 3.20, df = 8.70, p = 0.0113, two-tailed Welch’s t-test).

**Fig. 2.**
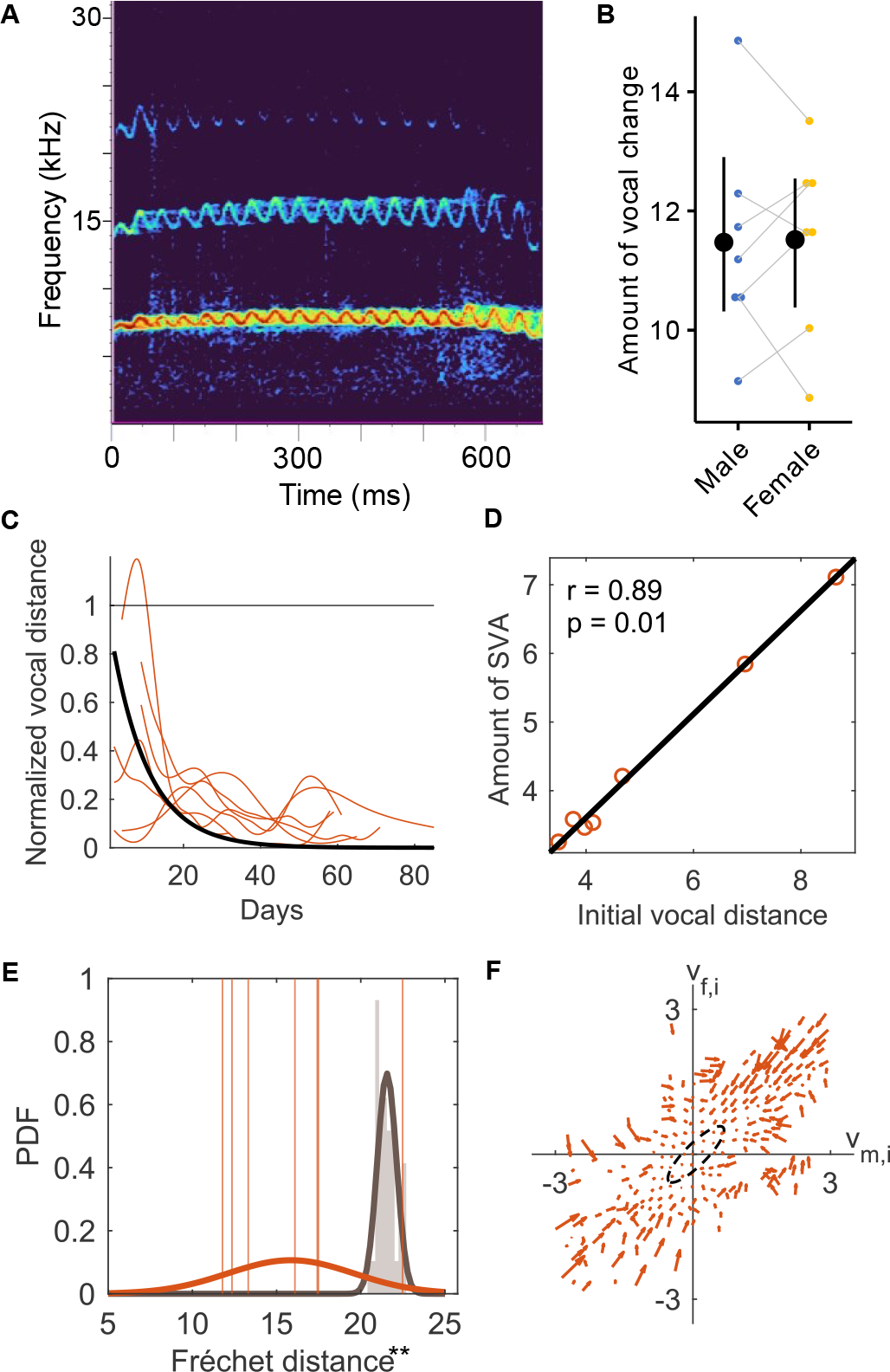
Properties of SVA in trills. (**A**) Example spectrogram of a trill call obtained using the Hann filter of size 512 samples, hop length of 256 samples, and DFT bin size of 512 samples. (**B**) Amount of vocal change undergone by males (n=7) and females (n=7) during SVA in trills. Each point is an individual. Male-female pairs are joined using grey lines. Black circles with error bars depict mean ± 95% bootstrapped CI. (**C**) Temporal progression of SVA in trills of n=7 pairs. Vocal distances are normalized to the before pairing condition. Green curves are spline fits for each pair, fit using MATLAB’s ‘smoothingspline’ with smoothing parameter of 0.1. Black curve depicts the first order exponential term fit to the population. (**D**) Correlation between the amount of SVA and Initial vocal distance for n = 7 pairs. ‘r’ is the Spearman’s correlation coefficient. (**E**) Dyadic acoustic feature synchrony in n=7 actual pairs (orange) compared to n=42 control pairings (grey) measured using Fréchet distance between the trajectories of the male and the female in a pair. Orange vertical lines depict the Fréchet distance values of actual pairs. Grey (normalized) histograms depict the distribution of Fréchet distance values of control pairs. Orange and grey curves are Gaussian probability distribution function fits to actual pairs and random pairs respectively. **p<0.01, two-tailed Welch’s t-test. (**F**) Phase portrait of normalized acoustic feature values of n = 7 pairs (see materials and methods for more details). The region approximately containing equilibrium points is marked with a dashed outline.

Finally, we visualized the dynamics of SVA in trills by plotting the acoustic feature values of females (v_f,i_) against their partner males (v_m,i_) on a phase portrait (Fig. 2F). We note 3 attributes of the phase portrait. (1) The equilibrium points lie approximately within an ellipse whose major axis lies along v_m,i_=v_f,i_. (2) Most arrows point toward the ellipse suggesting the system moves towards v_m,i_=v_f,i_. (3) Arrows further away from the ellipse are longer, representing the higher rate of change of feature values when the vocal distance between the male and the female was larger.

### Modelling the temporal dynamics of SVA in trills

The convergence of trills can be achieved in four ways. We developed a mathematical model for each and compared the properties of the models to the real marmoset SVA data. Each call was represented in the model using 20 acoustic features, and all models assumed that marmosets could modify each acoustic feature (v_m,i_ or v_f,i_, i^th^ acoustic feature of male or female calls respectively) at every time step (t) independently of the other. Each model was characterized using a separate learning rate parameter for the male (α_m_) and the female (α_f_), which, when put together, determined the rate of convergence of the pair during SVA. We also separately modelled errors E_m_X_t_ and E_f_X_t_ for males and females, respectively, where the errors are a product of the error term E and a Gaussian noise term X at time t (X ∼ N(0, 1)).

### Initial Auditory Template Matching (IATM)

In this model, marmosets are assumed to follow the following steps to achieve vocal convergence (Fig. 1A):

1. When paired, marmosets initially form an auditory template of the trills of their partner.
2. Marmosets modify their trills to match the auditory template.
3. Marmosets continue to modify their trills to match the template. The template itself is static and is not updated to account for changes in the partner’s trills.

IATM is the most parsimonious of all models as it does not require updating the auditory template. It is represented by the following equations:

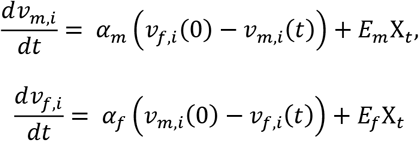

### Convergence to Intermediate Value (CIV)

In CIV, we assume that in addition to an acquired auditory template (from their partner), marmosets possess a template of their own trill (internal template), which might have previously played a role in vocal development. We build CIV as follows (Fig. 1B):

1. When paired, marmosets form an auditory template of the trills of their partner (acquired template) in addition to the internal template that they already possess.
2. Both the marmoset’s own internal template and the acquired template drive VPL. Marmosets modify their trills to match both templates.
3. Marmosets continue to modify their trills to match both templates. However, akin to IATM, the templates are not updated.

For the mathematical modelling of CIV, we chose the simplest case where both the internal and acquired templates equally affected the trills. The equations for CIV are as follows:

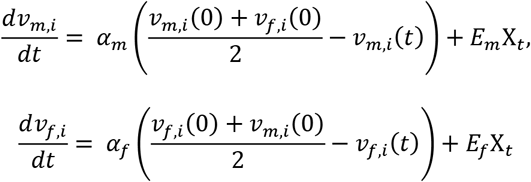

### Dynamic Auditory Template Matching (DATM)

Building on IATM, we add a layer of complexity by making the auditory template dynamic and sensitive to changes in the partner’s trills. This model can be summarized as follows (Fig. 1C):

1. When paired, marmosets form an auditory template of the trills of their partner.
2. Marmosets modify their trills to match the auditory template.
3. The template is continuously updated to account for changes in the partner’s trills, i.e., every time the trill structure of the partner changes, this changed trill acts as the new template. Marmosets modify their trills to match the updated template. Step 3 repeats.

In this model, marmosets will be able to account for errors or abrupt changes in the trills of the partner during SVA. DATM was modelled using the following equations:

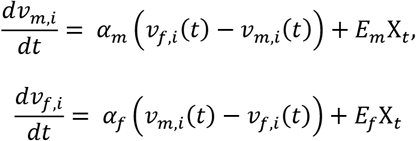

### Dynamic Convergence to Intermediate Value (DCIV)

Building upon CIV, we added a layer of complexity such that the internal and acquired templates are dynamic. According to DCIV (Fig. 1D):

1. Marmosets already have an internal template before pairing. When paired, marmosets additionally form the acquired template.
2. Trills are modified to match both templates.
3. Marmosets retain a memory of their own trill before modifying their trills. This becomes their new internal template. The acquired template is adjusted to account for the changes in the partner’s trills. Therefore, both the internal template and the acquired template are dynamic.

Similar to CIV, we chose the case where both the dynamic internal template and the dynamic acquired template are given equal weightage when marmosets match their trills to them. DCIV was modelled using the following equations:

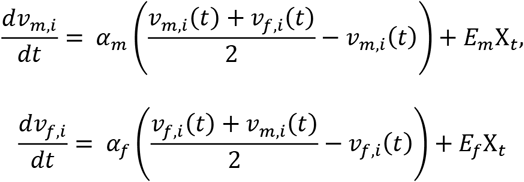

We can show that DCIV displays the same dynamics as DATM with half the learning rate as rearranging the above equation gives:

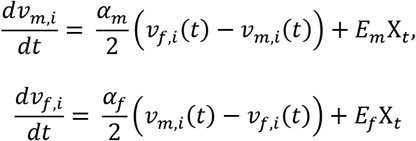

Therefore DATM/DCIV will be henceforth referred to as the “dynamic model”.

To be able to explain the synchronization of acoustic features between individuals in a pair, we added an error term to every model. Based on our observations of the nature of residuals during acoustic feature synchrony, where they decreased with time (Movie S1), we came up with 2 different error terms:

1. *Ε = f(t):* “Error is a function of time”. The classic learning curve where the error rate exponentially decreases with time given by:

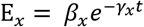

Where x=m for males and x = f for females. β and γ are error coefficients and t is time. To ensure that the dynamics shown by the model is invariant under the same error coefficients but linearly transformed initial acoustic feature values, the error term was modified as following:

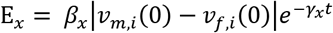
2. *E = f(dv/dt):* Arising from the possibility that the more the marmosets modify their calls, the more they are prone to making errors, we modelled the error to be a linear function of the amount of change in the acoustic feature value. This can be written as:

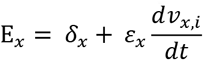

Where x=m for males and x=f for females. δ and ε are error coefficients, and t is time.

To make the model invariant under linear transformation of the initial acoustic feature values, we modified it to:

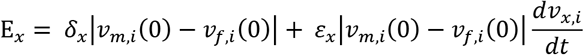

We simulated 7 virtual pairs with the trills of the virtual marmosets characterized by 20 acoustic features so that we could compare the temporal dynamics shown by the model to SVA in trills of actual pairs. We examined which of the models showed significant synchrony in acoustic features. For simplicity, we set the learning rates (α) and error coefficients (β, γ, δ and ε) of males and females to be equal. We varied these parameters within a suitable range (Table S2) and let the system evolve for 60 iterations (equivalent to 60 days). We found significant acoustic synchrony in a considerable range of parameter values in the case of the dynamic model with both types of error terms (Fig. 3E and 3H). In contrast, acoustic synchrony was only found in sparse patches for IATM and CIV only in the case of the *E = f(dv/dt)* error term. These results were robust to changes in acoustic feature initialization and the seed of the random number generator (Fig. S3 and S4).

**Fig. 3.**
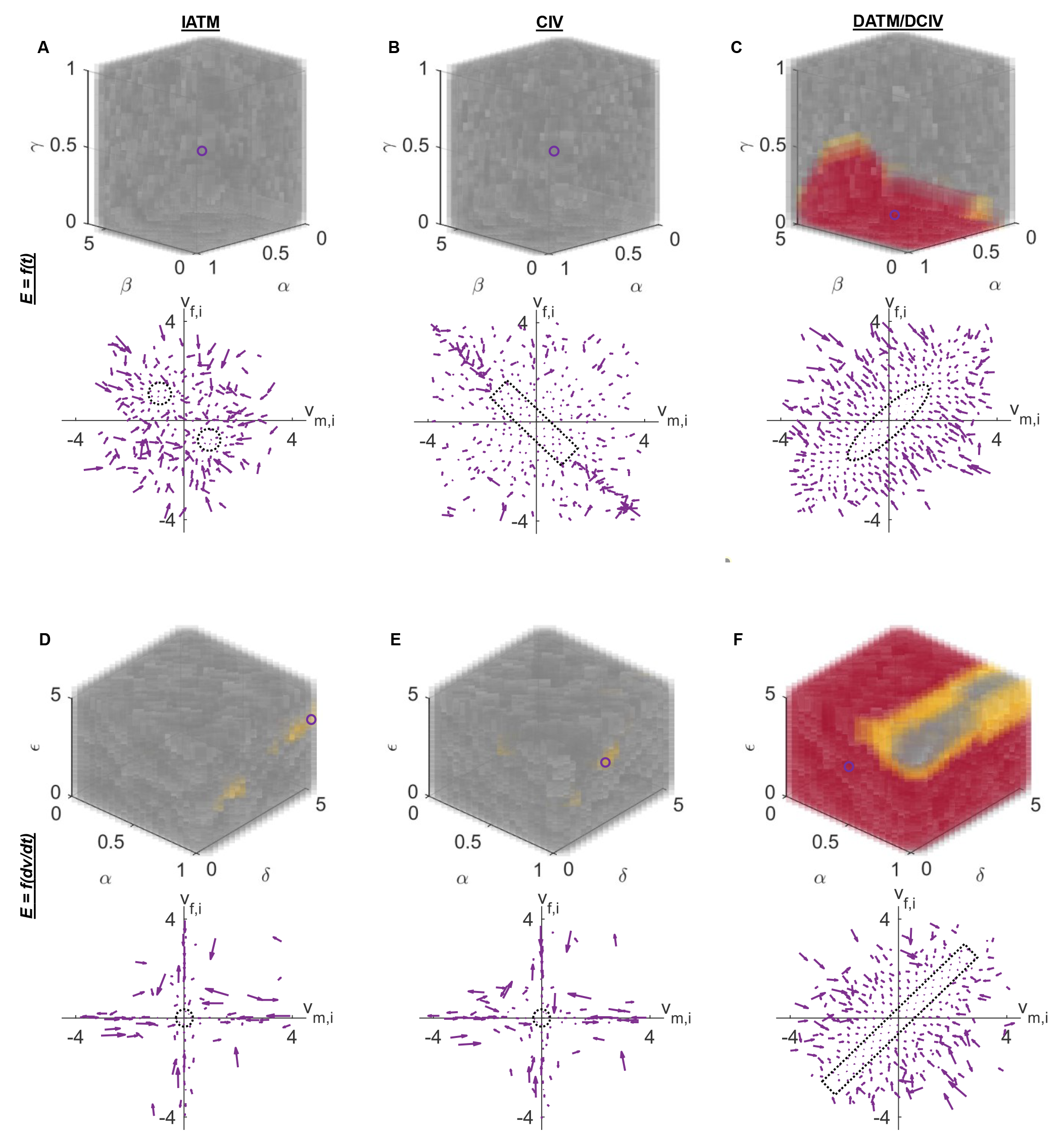
Behavior of the models. Top plots show the model parameter space. Model parameter values for which the acoustic features of the n=7 virtual pairs are significantly more synchronized than the n=42 virtual control pairs are marked in yellow if p<0.05, and in red if p<0.01 (two-tailed Welch’s t-test, acoustic trajectory similarity calculated using Fréchet distance). The rest of the points are filled in grey. The fill colors are made slightly transparent such that points behind them are visible. α is the learning rate while β, γ, δ and ε are error coefficients. Bottom plots depict the phase portrait for the model parameter values marked using open blue circles in top panels. The region of the phase portrait approximately containing equilibrium points is outlined with a black dotted curve. See materials and methods for more details regarding model simulations, synchrony calculations and phase portrait visualizations. Abbreviations: IATM = Initial Auditory Template Matching, CIV = Convergence to Intermediate Value, DATM = Dynamic Auditory Template Matching, DCIV = Dynamic Convergence to Intermediate Value.

Within the range of learning rate and error coefficient values for which significant acoustic synchrony was found, we randomly chose a set of values for the model, performed simulations for 60 iterations (equivalent to 60 days), and visualized the temporal dynamics on a phase portrait. The goal was to find the closest match to the phase portrait obtained from actual data (in Fig. 2F). Phase portraits of only the dynamic models were characterized by an approximately ellipsoid region of equilibrium points with the major axis along v_m,i_ = v_f,i_ (Fig. 3E and 3H). Even though the equilibrium points in the dynamic model with *E = f(dv/dt)* error term lie within a strip along v_m,i_ = v_f,i_ (Fig. 3F), we included the model for later analyses as it is possible that the strip is part of a larger ellipse. Based on the above observations, we concluded that the dynamic model best explains the dynamics of SVA in trills.

### Properties of the dynamic model

#### Bidirectional learning

With the selected model, we sought to determine the nature of the error term that would provide the best fit of the dynamic model to the data. The *E = f(t)* error term gave a better fit with significantly higher adjusted R^2^ values (Fig. 4A, n = 14, p = 1.2e-04, Wilcoxon signed-rank test) and substantially lower leave-one-out cross-validated normalized root mean square error (LOOCV NRMSE) values (Fig. 4B, n = 14, p = 1.2e-04, Wilcoxon signed-rank test). Further, the learning rates obtained from fitting the dynamic model to the trill data were similar between males and females (Fig. 4C, n = 7 pairs, t-value = -1.39, df = 6, p = 0.21, two-tailed paired t-test), confirming equal contributions of males and females towards vocal convergence during SVA.

**Fig. 4.**
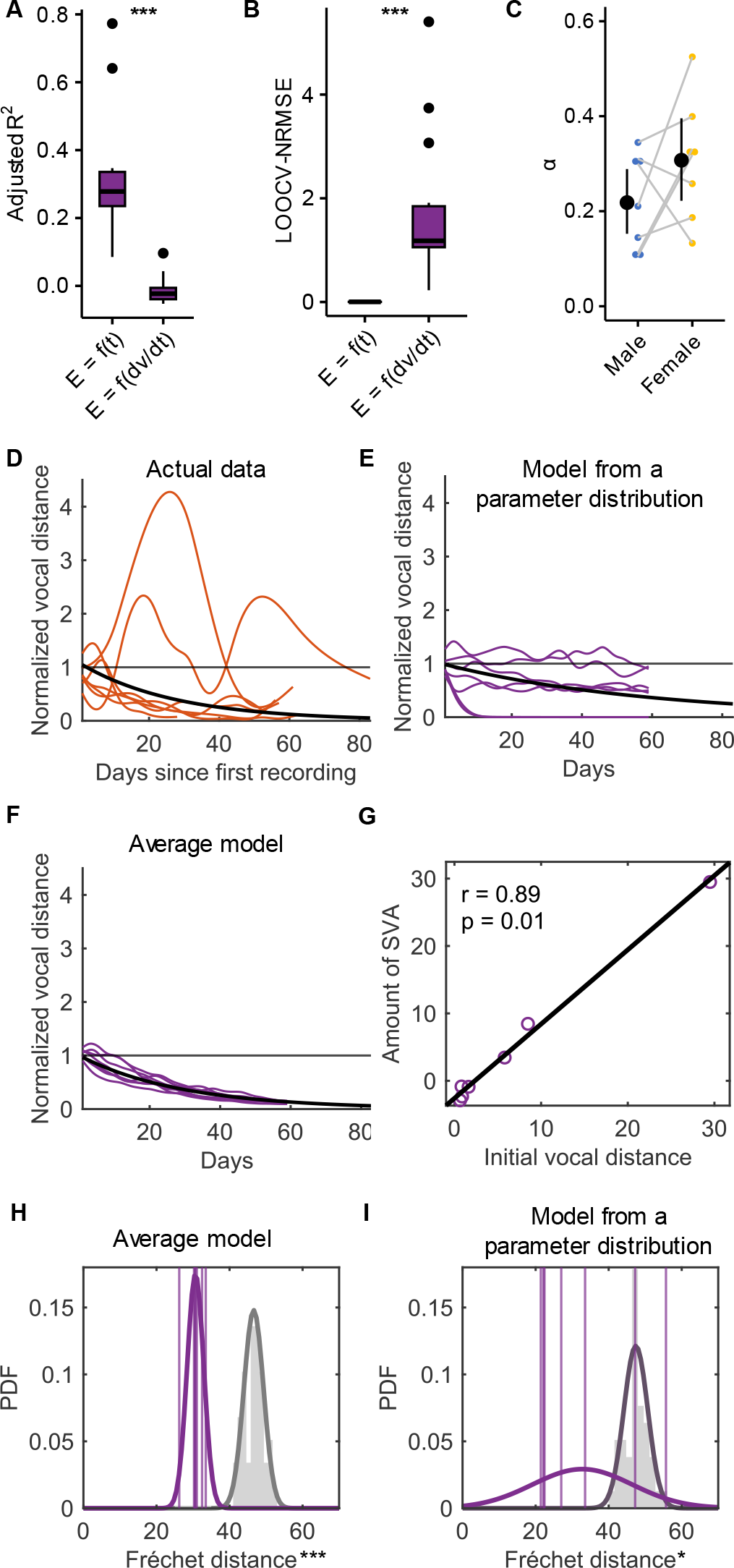
Properties of the dynamic model. (**A, B**) Box plots showing adjusted R^2^ (**A**) and leave-one-out cross-validated normalized root mean square error (LOOCV NRMSE) (**B**) of *E = f(t)* and *E = f(dv/dt)* fits to the residuals for n=14 individuals (***p<0.001, Wilcoxon signed-rank test). The residuals were obtained after a robust fit of the dynamic model without the error term to the data from each individual. (**C**) Learning rates (α) obtained after fitting the dynamic model to SVA data from each individual (n=14 individuals). Each point is an individual. Male-female pairs are joined using grey lines. Black circle with error bars depict mean ± 95% bootstrapped CI. (**D**) Temporal progression of SVA in actual data normalized to the vocal distance in the first recording after pairing. (**E, F**) Temporal progression of SVA in dynamic models with (**E**) parameters extracted from a distribution fit over actual parameters and (**F**) median of actual parameter values. (**G**) Correlation between the amount of SVA and initial vocal distance for n = 7 virtual pairs in the model. R is the Spearman’s correlation coefficient. (**H, I)** Acoustic feature synchrony in n=7 virtual pairs (purple) compared to n=42 virtual control pairings (grey) in the dynamic model simulated using (**H**) median of actual parameter values and (**I**) a distribution fit over actual parameter values, measured using Fréchet distance between the trajectories of the male and the female in a pair. Purple vertical lines depict the Fréchet distance values of virtual pairs. Grey (normalized) histograms depict the distribution of Fréchet distance values of virtual control pairs. Purple and grey curves are Gaussian probability distribution function fits to actual pairs and random pairs respectively. *p<0.5, ***p<0.001, two-tailed Welch’s t-test.

#### Exponential decrease of vocal distance with time

We fit a distribution over the learning rates (Gaussian) and error coefficients (gamma) from the dynamic model fit separately for males and females. From these distributions, we randomly extracted learning rates and error coefficients for the simulation. We performed simulations of 7 models (each symbolizing one actual marmoset pair in the data) for 60 iterations (equivalent to 60 days) and compared the temporal progression of SVA in the model to actual data. We repeated the same but this time using the median learning rates and error coefficients and simulated the “average model”. The nature of the decrease in vocal distance between virtual pairs in the models closely resembled the actual ones, both showing exponential decay (Fig. 4D, E and F).

#### Sensitivity to initial vocal distance

Along with studying the temporal progression of SVA in the dynamic models, we also examined the relationship between the total amount of SVA in a pair between t = 0 and t = 60 and the initial vocal distance. We found the amount of SVA in the dynamic models to be positively correlated with the initial vocal distance (Fig. 4G, Spearman’s correlation coefficient r = 0.89, p = 0.01).

#### Dyadic acoustic feature synchrony

Even though our model selection was based on the presence of acoustic feature synchrony for a wide range of model parameter values, the parameter values of males and females were tied to be equal during model simulations for Fig. 3A-2F. We sought to confirm whether acoustic feature synchrony remained in the simulated models. We found significant vocal synchrony in the 7 virtual pairs compared to virtual control pairs, confirming that acoustic feature synchrony can arise automatically in such models (Fig. 4H for median parameters, t-value = -16.65, df = 9.05, p = 4.28e-08, two-tailed Welch’s t-test; Fig. 4I for parameters taken from a distribution fit, t-value = -2.827, df = 6.117, p = 0.030, two-tailed Welch’s t-test). We visualized this acoustic synchrony by plotting the trajectories taken by the virtual male and the female in the acoustic feature space upon dimensional reduction (Movie S2).

## Discussion

### The properties of vocal learning in adult marmosets

When marmosets were paired with a new partner, their calls became more similar to each other. This accommodation was strongest in trills, which are close-distance contact calls^18^. Using highly accurate representations of trill calls^21^, we found four distinct patterns when pairs of marmosets engage in SVA. First, we were interested in who would accommodate to whom (i.e., the directionality). One hypothesis is that individuals more interested in forming a pair bond would accommodate more than their partner (10). This could arguably be the males because they are interested in siring offspring as soon as possible and also tend to show higher rates of affiliative behaviors early during pair formation^27^, or females, because as cooperative breeders, they are interested in forming a strong bond to ascertain future paternal contributions to infant care^28^. We neither found that males would vocally accommodate more to females nor vice versa, but both partners adjusted their own calls to approximately equal amounts (Fig. 2B). This is in line with the high levels of mutual affiliative behaviors in newly formed marmoset pairs^27^ and a mutual interest in a strong bond since both partners mutually depend on a durable, lasting bond to successfully reproduce^28^.

Second, the decrease in vocal distance is a gradual, exponential process (Fig. 2C). This contrasts with developmental changes in both human and marmoset infants, which are characterized by sudden transitions in the acoustic properties of vocalizations^29^. In contrast, our investigation of acoustic changes in adult marmosets suggests a gradual temporal progression with an exponential decrease in vocal distance within dyads. Such a gradual progression while matching a template is consistent with studies on pitch recovery in adult zebra finches^30^ and indications of gradual temporal progression of SVA in humans^31^.

Third, the total amount of SVA undergone by a pair is positively correlated to the initial vocal distance between the pair (Fig. 2D). This is logically coherent due to the fact that a large initial vocal distance provides more room for a pair to converge in the vocal space. It is in line with a previous finding on a smaller dataset of marmoset trills using spectral features^19^ and suggests that VPL is sensitive to initial conditions that must be taken into account when comparing across pairs and developing models of VPL.

The fourth and most prominent pattern in marmoset SVA was that the calls of dyads show synchronized changes in acoustic features over time, i.e., they move together through the acoustic space (Fig. 2E, Movie S1). Importantly, synchronization was higher in real dyads compared to all virtual control pairs composed of all possible different-sex dyads in the data set. These findings are consistent with previous observations based on traditional acoustic analyses^22^. Intriguingly, the tendency to synchronize has also been reported in other domains in marmosets, such as in oxytocin profiles of strongly bonded marmosets^32^ or in behavioral alignment^33,34^ and thus, may well be part of a general pattern of bio-behavioral synchrony^35^.

### The dynamic model best describes adult marmoset vocal learning

The four observed properties of adult marmoset SVA were best instantiated by the dynamic model (Fig. 3C, Fig. 4C-I). The model has implications for understanding VPL in adult marmosets. In particular, it predicts the presence of mechanisms responsible for the formation of the dynamic auditory template and matching of trills to the template. Neural mechanisms for detecting differences between the partner’s trills and the marmoset’s own trills may lie in the auditory cortex. Neurons in the marmoset auditory cortex have been shown to be modulated in two different ways during self-initiated vocalizations; one population of neurons is excited during vocal production while the other is suppressed, with the latter being greater in number^36,37^. We hypothesize that excited neurons tuned to the marmoset’s own trills and the suppressed neurons tuned to the partner’s trills may together play a role in detecting differences between the structure of one’s own trill and the partner’s trill. Moreover, neurons in the marmoset auditory cortex suppressed by self-generated vocalizations have been shown to be tuned to particular values of frequency and other acoustic features of auditory stimuli, where stimuli included vocalizations of other marmosets in the colony^38^. Such neurons, along with a downstream circuit, may play a role in the dynamicity of the auditory template. A “comparison circuit” capable of detecting differences between the responses of the excited neurons and neurons encoding the dynamic template can drive marmosets to update their trills to match their partners’, ultimately leading to SVA.

The dynamic model suggests that synchronization in acoustic parameters during SVA can automatically arise when marmosets account for changes in their partner’s trills and modify their own trills accordingly. The model, therefore, provides direct evidence for the Interactive Alignment Model (IAM)^39–41^ proposed for vocal accommodation in humans. IAM states that SVA arises automatically due to a tight link between speech production and perception in human interlocutors. Importantly, this does not exclude additional effects of social bond strength on the amount of SVA, as suggested by the Communication Accommodation Theory (CAT)^42,43^. This is because the dynamic model cannot explain the variability in learning rates across individuals. We propose that while the mechanisms of SVA are fundamental in driving vocal convergence in adult marmosets and humans, social factors contribute to fine-tuning of the phenomena, which in extreme cases may also lead to vocal divergence as seen during negative social interactions in humans^44^. Future studies on the biological function of SVA, perhaps in particular of effects of social bond strengths on the amount of SVA in marmosets, will help disentangle the contributions of fundamental, possibly innate mechanisms and social factors towards SVA.

A limitation of the dynamic model is that it assumes that marmosets can vary their acoustic features independently of each other. This may not always be true, as the physiology of the vocal apparatus may add restrictions to how much a particular acoustic feature can be modified given the changes in other acoustic features. Detailed study of the structure of the marmoset vocal apparatus and associated organs, along with computational modelling of vocal production, will help refine the dynamic model.

Together, the above evidence gives way to the intriguing possibility that divergent mechanisms can underlie VPL in infant and adult animals. First, the transitions in the acoustic properties are gradual in adults but abrupt in immatures. Second, since infant VPL is directional with a stable set of target (adult) vocalizations, an updating of the auditory template is not necessary. Our model suggests that VPL in adult marmosets requires constant updating of the auditory template to account for changes in partners’ vocalizations. Importantly, to further explore this possibility, future studies will have to carefully compare different types of VPL (e.g., adjusting of acoustic structure as in SVA, acquisition of novel vocalizations and words, acquisition of call combinations and syntactic rules) in both immatures and adults of the same species to unveil which mechanisms are similar/dissimilar during immature vs adult vocal learning. Given the need for solid comparative evidence to understand the evolutionary origin of human language, such systematic comparisons are an effortful but very much needed next step.

## Methods

### Experimental Subjects

20 common marmosets (*Callithrix jacchus*), 10 males and 10 females were used to obtain vocal recordings. Marmosets were housed in enclosures (1.8m x 2.4 m x 3.6m) along with at least one other group member. Each group had access to their own outdoor enclosure (1.8m x 2.4m x 4.6m) and a common experimental room. Marmosets were fed a predetermined amount of mush in the morning, cooked and raw vegetables during noon, and insects, cottage cheese or gum Arabic as snacks after 12:00 hours local time. Ad libitum access to water was provided at all times. All experiments were approved by Zürich’s cantonal veterinary office (license ZH223/16).

### Pair formation and vocal recordings

All experiments and vocal recordings were performed by Zürcher et al.^18^. Marmoset vocalizations were recorded under two conditions: before pairing and after pairing. During the before pairing condition, individuals were recorded in their home enclosure on multiple days spread over two to three weeks. Each recording session being about 30 minutes long. For the after pairing condition, males and females were paired (each of the 10 males paired with a female), and vocalizations were recorded at regular intervals on about 15 days (15 ± 4 days, median ± std.) in the period starting from the day of pair formation to up to 60 days after pairing (61 ± 16 days, median ± std.). Individuals were recorded in their home enclosure or the experimental room.

Marmoset calls were obtained using a condenser Microphone (CM16/CMPA, Avisoft Bioacoustics, Germany) connected to Avisoft UltraSoundGate 116H (Avisoft Bioacoustics, Germany) at a sampling rate of 62500 Hz and source (individual) identity of all calls were simultaneously labeled using Avisoft Recorder (Avisoft Bioacoustics, Germany). Calls were visualized and segmented using Avisoft Pro (Avisoft Bioacoustics, Germany), and each call was saved in a separate file. See Zürcher et al.^18^ for detailed information about pair formation, recording procedure, and processing.

### Properties of SVA in trills

To study the Properties of SVA, we chose 7 pairs (out of the 10) for which recordings with high temporal resolution were available (7 pairs: 7 males and 7 females). For acoustic feature extraction from animal vocalizations, Highly Comparative Time Series Analysis (HCTSA^45,46^) has been shown to be an effective method^21,47^. This method extracts numerous features directly from the raw acoustic waveform (pressure time series). We extracted features from 5842 marmoset trills recorded both before pairing (1088) and after pairing (4754) of individuals using HCTSA.

Previous studies have shown features from animal vocalizations extracted by HCTSA to be useful for supervised classification based on context and source properties^21,47^. In particular, tree-based boosting methods can classify marmoset calls based on the sex and identity of the source with high accuracies^21^. Such classifiers also identify the acoustic features in which individual identity signatures are most likely to lie. The top 20 most important features (out of 3255) for classifying marmoset trills were determined. For exact details of the classifier, and classification accuracies, see Phaniraj et al. ^21^. For a list of the top 20 features and their description, see Table S1.

Using the top 20 feature set, we aimed to quantify vocal distances between the trills of individuals of every pair during SVA. For this, we calculated the Euclidean distance between centroids wherein the position of the centroids of the two clusters (one for each individual in a pair) in the 20-dimensional feature space (top 20 feature space) was determined, and the Euclidean distance between them was calculated.

To compare the amount of vocal change undergone by males and females during SVA, we selected the last day after pairing on which both individuals in a pair were recorded. Features were normalized (z-scored), and the distance between the calls of an individual on the last day and before that individual was paired with a partner was calculated. We used two-tailed paired t-tests to detect significant differences in the amount of vocal change undergone by males and females. To visualize the temporal progression of SVA, we first calculated the vocal distance between individuals using normalized features before they were paired to set a baseline. We then calculated the vocal distance between pairs on all days after pairing when both individuals were recorded. To determine the relationship between the amount of SVA and initial vocal distance, the amount of decrease in vocal distance between a pair from the first day of recording after pairing and the last day of recording was correlated with the vocal distance between the pair on the first day of recording after pairing.

To test for the presence of synchrony in vocal features during SVA, we first determined the centroid of the cluster (in the normalized feature space) of all calls given by an individual of a pair on the days they were recorded after pairing with a partner. Next, we performed linear interpolation and obtained centroid locations on the days with no recordings. This provided us with “vocal trajectories” of pairs when centroids of all days were connected. However, the amount of convergence between pairs would influence how similar the vocal trajectories of the individuals in the pair are. We, therefore, decomposed the vocal trajectories into a linear trend and the residuals to remove this influence. For this, multivariate regression was performed on the vocal trajectory (with day as the predictor variable), and residuals were obtained. To quantify vocal feature synchrony, we determined the trajectory similarity between residuals obtained for an individual and its partner. We used two measures for this:

1. *Dynamic Time Warping (DTW) distance*: The DTW implementation on multivariate time series data^48^ with a window parameter of 1 (maximum window size) was used to calculate the warping cost, which was reported as the distance. Large distance values represent low trajectory similarity values.
2. *Fréchet distance*: The discrete Fréchet distance (MATLAB file exchange #31922) between the two trajectories was calculated. Larger values indicate lower similarity.

As a control, we also calculated trajectory similarity between the residuals of all possible non-bonded male-female pairings in the data. For N pairs, N actual similarity values and N(N-1) control similarity values could be calculated. We used two-tailed Welch’s t-tests for samples with unequal variances to detect significant differences in the Fréchet and DTW distance measures of actual pairs and control pairs. Additionally, we repeated the above analyses without controlling for the amount of convergence between pairs.

Dyadic acoustic feature synchrony could also be measured using traditional spectral features. To test if traditional analyses would also detect synchrony, we extracted four commonly used spectral features for marmoset acoustic analyses^29,49^ from all calls: mean fundamental frequency, mean spectral entropy, frequency of amplitude modulation and call duration. For this, all calls were filtered to be between the marmoset trill frequency range (4 – 10kHz). The MATLAB function ‘pitch’ was used to estimate the fundamental frequency, and its average over the entire call was used. For determining mean spectral entropy, the MATLAB function ‘spectralEntropy’ was used, and its average over the entire call was reported. To estimate the frequency of amplitude modulation, Hilbert transform was applied (using MATLAB function ‘hilbert’) to the waveform to first obtain the amplitude envelope and its frequency was determined. We performed Principal Component Analysis over z-scores of these four features and determined the Principal Acoustic Component (PAC). Then, from the time series of PAC values for individuals in a pair, we calculated the mean phase coherence. For this, we first bandpass filtered the two time series from a pair between the values [(0.125/length of time series) to 0.5 cycles per day] to remove both extremely low and extremely high frequencies that could contribute to these fluctuations, which are not of our interest. After filtering, we determined instantaneous phases by performing a Hilbert transform using the MATLAB function ‘hilbert’. Finally, we calculated the mean phase coherence using the formula:

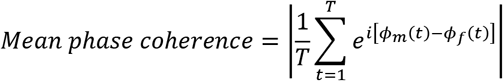

Where ϕ_*m*_(*t*) and ϕ_*f*_(*t*) are the instantaneous phases of the male’s and female’s PAC at instant t and T is the duration (days) between day of pairing and the last day of recording after pairing. The above analysis was also repeated with fundamental frequency alone, onto which PAC was heavily loaded.

We visualized the vocal trajectories taken during SVA of all pairs using a phase portrait. For a given day after pairing and a given pair, we determined the centroid of all calls in the normalized feature space for the male and the female separately. For every acoustic feature i, we plotted the value for the female (v_f,i_ – y coordinate) against the value for the male (v_m,i_ – x coordinate). Then, we determined the rate of change of the acoustic feature between that day and the next day of recording (ratio of the difference in acoustic feature value to the number of days passed) for both the male (dv_m,i_/dt) and the female (dv_f,i_/dt). Arrows originating from the points (v_m,i_, v_f,i_) with directional components (dv_m,i_/dt, dv_f,i_/dt) were plotted. We repeated this for all pairs and all days of recording after pairing. To prevent overcrowding of arrows in the phase portrait, we further chose a range of v_m,i_ and v_f,i_ symmetric around 0 that contained the majority of the data. Within this range, we divided the entire plot into 625 (25x25) squares. Within each square, the mean of all arrow vectors was calculated and the entire box was represented by 1 mean arrow vector.

### Modelling the temporal dynamics of SVA in trills

Our goal was to come up with the simplest set of mathematical equations that could show all the properties of the temporal dynamics of SVA in trills. Inspired by the auditory template hypothesis of vocal learning in songbirds^25,26^, we hypothesized four ways marmoset pairs could achieve convergence in trills and developed mathematical equations for modelling them. Additionally, to be able to explain the synchronization of acoustic features between individuals in a pair, based on our observations of the nature of residuals during acoustic feature synchrony, we added two types of error terms to every model. (1) As the classical learning curve, the error rate (Ε) decreases exponentially with time (“Error is a function of time”, *E = f(t)*), or (2) The more the individuals change the acoustic features of trills, the more prone they are to making errors (“Error is the function of the rate of change of acoustic feature value”, *E = f(dv/dt)*). Both the error terms are comprised of 2 error coefficients, β and γ for *E = f(t)*, and δ and ε for *E = f(dv/dt)*. Virtual males and females were given error coefficient values separately. The error terms were constructed such that the model is invariant under changes in the linear transformation of initial conditions. The models are represented using the following equations:

1. *Initial Auditory Template Matching (IATM) with E = f(t) error term:*

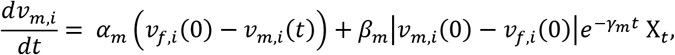

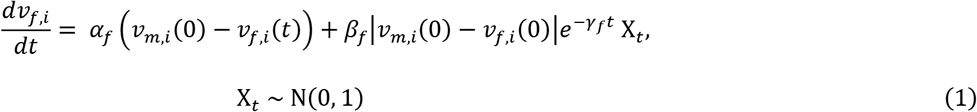
2. *Initial Auditory Template Matching (IATM) with E = f(dv/dt) error term:*

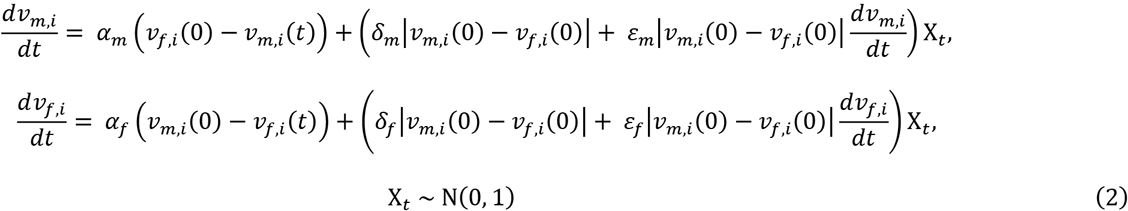
3. *Convergence to Intermediate Value (CIV) with E = f(t) error term:*

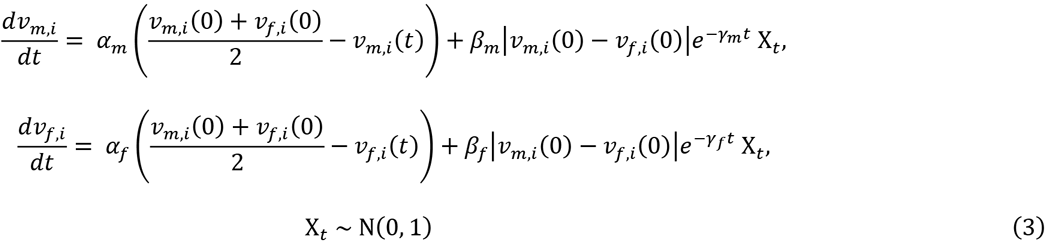
4. *Convergence to Intermediate Value (CIV) with E = f(dv/dt) error term:*

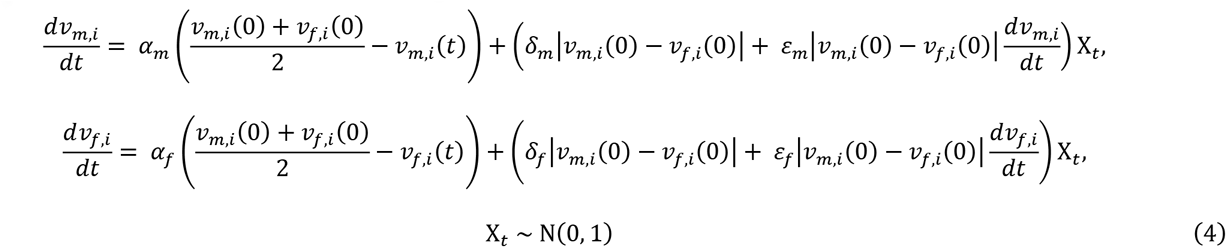
5. *Dynamic Auditory Template Matching (DATM) with E = f(t) error term:*

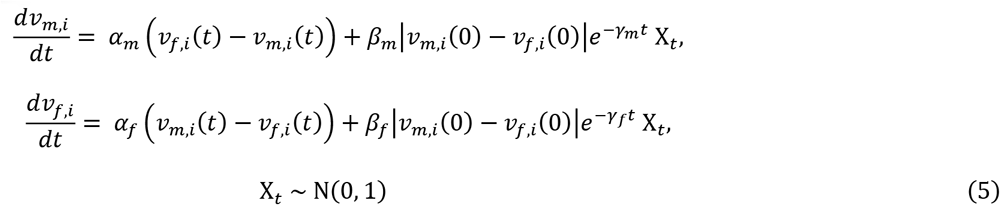
6. *Dynamic Auditory Template Matching (DATM) with E = f(dv/dt) error term:*

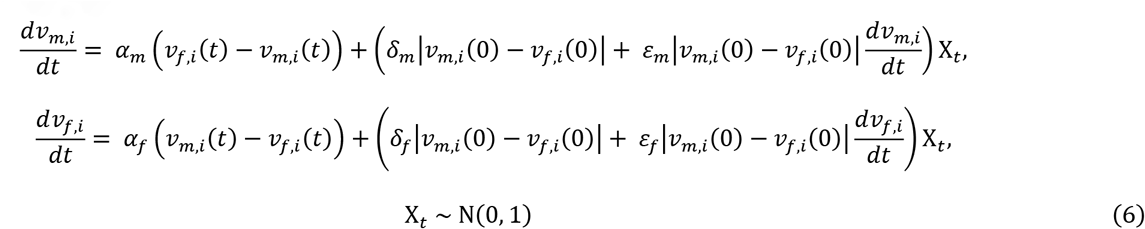
7. *Dynamic Convergence to Intermediate Value (DCIV) with E = f(t) error term:*

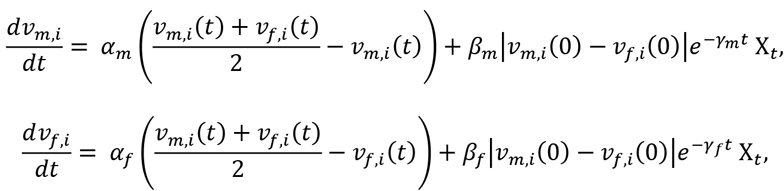

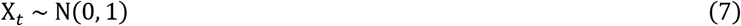
8. *Dynamic Convergence to Intermediate Value (DCIV) with E = f(dv/dt) error term:*

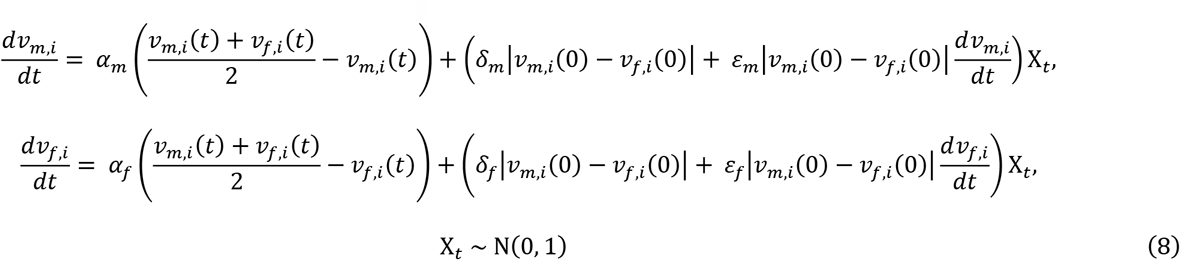

In all the above models, v_m,i_ and v_f,i_ are the values of the i^th^ acoustic feature of male and female calls respectively, t is time, α_m_ and α_f_ are the learning rates of the male and the female respectively, β, γ, δ and ε are error coefficients, and *X*_*t*_ is Gaussian noise at time t. The Error term was further modelled in 2 different ways:

We simulated 7 virtual marmoset pairs and 20 acoustic feature values (same as in the actual data). We performed numerical simulations of equations (1-8) using Euler’s method and the parameter value range mentioned in Table S2. Gaussian noise was modelled using MATLAB’s ‘randn’ function, and acoustic feature values were initialized from a uniform distribution using MATLAB’s ‘rand’ function. Then, for each simulation, we tested for the presence of synchrony using the Fréchet distance measure, as described in the previous section. We compared the Fréchet distance between the trajectories (of the residuals of the acoustic feature values) of the virtual marmoset pairs with the virtual control pairs (similar to the case with original data, for 7 virtual pairs, 7 actual similarity values and 42 (7x(7-1)) control similarity values could be calculated). Fréchet distance was preferred over DTW distance as calculating DTW distance requires setting a window parameter which affects the output. Fréchet distance does not require setting any such parameter. Later, we examined if the pattern of synchronization was robust to changes in initial acoustic parameter values or the seed of the random number generators.

Finally, for each model, we selected a set of parameters for which synchronization was present, i.e., the Fréchet distance values for the 7 virtual pairs were significantly lower than the 42 (7x(7-1)) virtual control pairs. Then for those parameters, we visualized the temporal dynamics of the model by plotting a phase portrait. The steps followed are the same as those for the original data described in the previous section.

### Temporal dynamics of the best model

We studied the temporal dynamics of the models in which synchrony was present under a significant range of parameter values and whose phase portraits were qualitatively similar to the phase portrait obtained using actual marmoset data. First, we fit the model without the Gaussian noise term to the actual data using MATLAB’s ‘robustfit’ with the ‘cauchy’ weight function and its default tuning constant and obtained the learning rates (α_m_ or α_f_) for every individual. The ‘cauchy’ weight function was chosen as it is smooth, differentiable, and does not completely exclude any outliers. Then, we inspected which kind of error term fit the residuals the best. For this, we discretized dv/dt values and binned all residuals into the relevant t (day) or dv/dt. The bin size for the *Ε = f(t)* model was 1 day, whereas the ideal bin size for the *Ε = f(dv/dt)* model was determined using the Freedman-Diaconis rule. We fit a Gaussian distribution on the residuals within each bin and obtained the standard deviation of the distribution for that bin. We then checked whether the standard deviation values followed *E = f(t)* error type or *E = f(dv/dt)*. To do so, we compared the adjusted R-square values and leave-one-out cross-validated normalized root mean square error (LOOCV-NRMSE) values of both the error equation fits. Wilcoxon signed-rank tests were used to detect significance during comparisons. The model with the higher adjusted R-square and lower LOOCV-NRMSE was chosen as the better error model.

For the virtual males in the best model overall, we first set the learning rate and error coefficients to the median of the values given by the model when fit on data from actual males. The same was done to obtain the coefficients for the virtual females. We studied the temporal progression of SVA of the best model by plotting the vocal distance between the virtual males and the virtual females in the model against the number of days. Here too, we simulated as many virtual marmoset pairs and acoustic parameter values as there were in the actual data (7). We compared the temporal progression plot to a similar plot on the data obtained from actual marmoset pairs. Then, for the particular coefficient values chosen for the best model, we checked whether the virtual pairs in the model were significantly more synchronized than control pairs using the procedure described in the previous sections for testing the presence of dyadic acoustic synchrony. Finally, we repeated this analysis, but this time by fitting a distribution on the parameters instead of taking the median. We fit a distribution over the learning rates (Gaussian) and error coefficients (gamma) from the dynamic model fit separately for males and females. From these distributions, we randomly extracted learning rates and error coefficients and performed the above-mentioned simulations.

## Acknowledgements

We are thankful to Yvonne Zürcher for the marmoset vocal accommodation dataset and to Raghav Rajan and Richard Hahnloser for discussions at various stages of the project. This project was supported by the Swiss National Science Foundation (grant number 31003A_149796, the NCCR Evolving Language, agreement number 51NF40_180888) and the European Research Council (ERC) under the European Union’s Horizon 2020 research and innovation programme (grant agreement No. 101001295). N.P. was a recipient of a grant from the A. H. Schultz Foundation.

## Author contributions

N.P., K.W. and J.M.B developed the concept and designed the study. N.P. developed the methods and models with inputs from J.M.B.

N.P performed the analyses with inputs from K.W and J.M.B. N.P wrote the original draft. K.W. and J.M.B reviewed and edited the draft and provided supervision throughout the project. J.M.B acquired funding and administered the project.

## Supplementary information

Supplementary information is available for this paper.

## References

1. Janik, V. M. & Slater, P. J. B. The different roles of social learning in vocal communication. Animal Behaviour 60, 1–11 (2000).

2. Janik, V. M. & Knörnschild, M. Vocal production learning in mammals revisited. Philosophical Transactions of the Royal Society B: Biological Sciences 376, 20200244 (2021).

3. Vernes, S. C. et al. The multi-dimensional nature of vocal learning. Philosophical Transactions of the Royal Society B: Biological Sciences 376, 20200236 (2021).

4. The Neuroethology of Birdsong. vol. 71 (Springer International Publishing, 2020).

5. Searcy, W. A., Soha, J., Peters, S. & Nowicki, S. Variation in vocal production learning across songbirds. Philosophical Transactions of the Royal Society B: Biological Sciences 376, 20200257 (2021).

6. Soha, J. The auditory template hypothesis: a review and comparative perspective. Animal Behaviour 124, 247–254 (2017).

7. Marler, P. A comparative approach to vocal learning: song development in white-crowned sparrows. Journal of comparative and physiological psychology 71, 1 (1970).

8. Ikeda, M. Z., Trusel, M. & Roberts, T. F. Memory circuits for vocal imitation. Current Opinion in Neurobiology 60, 37–46 (2020).

9. Bolhuis, J. J. & Moorman, S. Birdsong memory and the brain: in search of the template. Neuroscience & Biobehavioral Reviews 50, 41–55 (2015).

10. Ruch, H., Zürcher, Y. & Burkart, J. M. The function and mechanism of vocal accommodation in humans and other primates. Biological Reviews 93, 996–1013 (2018).

11. Pardo, J. S., Pellegrino, E., Dellwo, V. & Möbius, B. Special issue: Vocal accommodation in speech communication. Journal of Phonetics 95, 101196 (2022).

12. Bernhold, Q. S. & Giles, H. Vocal Accommodation and Mimicry. J Nonverbal Behav 44, 41–62 (2020).

13. Miller, C. T. et al. Marmosets: a neuroscientific model of human social behavior. Neuron 90, 219–233 (2016).

14. Burkart, J. M. & Finkenwirth, C. Marmosets as model species in neuroscience and evolutionary anthropology. Neuroscience Research 93, 8–19 (2015).

15. Burkart, J. M. et al. A convergent interaction engine: vocal communication among marmoset monkeys. Philosophical Transactions of the Royal Society B: Biological Sciences 377, 20210098 (2022).

16. Eliades, S. J. & Miller, C. T. Marmoset vocal communication: behavior and neurobiology. Developmental neurobiology 77, 286–299 (2017).

17. Zürcher, Y. & Burkart, J. M. Evidence for dialects in three captive populations of common marmosets (Callithrix jacchus). International Journal of Primatology 38, 780–793 (2017).

18. Zürcher, Y., Willems, E. P. & Burkart, J. M. Are dialects socially learned in marmoset monkeys? Evidence from translocation experiments. PLOS ONE 14, e0222486 (2019).

19. Zürcher, Y., Willems, E. P. & Burkart, J. M. Trade-offs between vocal accommodation and individual recognisability in common marmoset vocalizations. Scientific Reports 11, 1–10 (2021).

20. Priva, U. C. & Sanker, C. Limitations of difference-in-difference for measuring convergence. Laboratory Phonology 10, (2019).

21. Phaniraj, N., Wierucka, K., Zürcher, Y. & Burkart, J. M. Optimising source identification from marmoset vocalizations with hierarchical machine learning classifiers. 2022.11.19.517179 Preprint at 10.1101/2022.11.19.517179 (2022).

22. Zürcher, Y. Vocal learning and flexibility in the communication of common marmosets (Callithrix jacchus). (University of Zurich, 2020).

23. Fréchet, M. M. Sur quelques points du calcul fonctionnel. Rendiconti del Circolo Matematico di Palermo (1884-1940) 22, 1–72 (1906).

24. Sakoe, H. & Chiba, S. Dynamic programming algorithm optimization for spoken word recognition. IEEE transactions on acoustics, speech, and signal processing 26, 43–49 (1978).

25. Konishi, M. The Role of Auditory Feedback in the Control of Vocalization in the White-Crowned Sparrow1. Zeitschrift für Tierpsychologie 22, 770–783 (1965).

26. Marler, P. & Nelson, D. Neuroselection and song learning in birds: species universals in a culturally transmitted behavior. Seminars in Neuroscience 4, 415–423 (1992).

27. Evans, S. & Poole, T. B. Long-term changes and maintenance of the pair-bond in common marmosets, Callithrix jacchus jacchus. Folia primatologica 42, 33–41 (1984).

28. Finkenwirth, C. & Burkart, J. M. Why help? Relationship quality, not strategic grooming predicts infant-care in group-living marmosets. Physiology & behavior 193, 108–116 (2018).

29. Varella, T. T., Zhang, Y. S., Takahashi, D. Y. & Ghazanfar, A. A. A mechanism for punctuating equilibria during mammalian vocal development. PLOS Computational Biology 18, e1010173 (2022).

30. Zai, A. T., Stepien, A. E., Giret, N. & Hahnloser, R. H. R. Goal-directed vocal planning in a songbird. 2022.09.27.509747 Preprint at 10.1101/2022.09.27.509747 (2023).

31. Tobin, S. J. Effects of native language and habituation in phonetic accommodation. Journal of Phonetics 93, 101148 (2022).

32. Finkenwirth, C., van Schaik, C., Ziegler, T. E. & Burkart, J. M. Strongly bonded family members in common marmosets show synchronized fluctuations in oxytocin. Physiology & Behavior 151, 246–251 (2015).

33. Koski, S. E. & Burkart, J. M. Common marmosets show social plasticity and group-level similarity in personality. Scientific reports 5, 8878 (2015).

34. Brügger, R. K., Willems, E. P. & Burkart, J. M. Looking out for each other: Coordination and turn taking in common marmoset vigilance. Animal Behaviour 196, 183–199 (2023).

35. Feldman, R. The neurobiology of human attachments. Trends in cognitive sciences 21, 80–99 (2017).

36. Eliades, S. J. & Wang, X. Sensory-motor interaction in the primate auditory cortex during self-initiated vocalizations. Journal of neurophysiology 89, 2194–2207 (2003).

37. Eliades, S. J. & Wang, X. Comparison of auditory-vocal interactions across multiple types of vocalizations in marmoset auditory cortex. Journal of Neurophysiology 109, 1638–1657 (2013).

38. Eliades, S. J. & Wang, X. Contributions of sensory tuning to auditory-vocal interactions in marmoset auditory cortex. Hearing Research 348, 98–111 (2017).

39. Pickering, M. J. & Garrod, S. The interactive-alignment model: Developments and refinements. Behavioral and Brain Sciences 27, 212–225 (2004).

40. Pickering, M. J. & Garrod, S. An integrated theory of language production and comprehension. Behavioral and brain sciences 36, 329–347 (2013).

41. Pickering, M. J. & Garrod, S. Understanding dialogue: Language use and social interaction. (Cambridge University Press, 2021).

42. Giles, H., Coupland, N. & Coupland, I. 1. Accommodation theory: Communication, context, and. Contexts of accommodation: Developments in applied sociolinguistics 1, (1991).

43. Giles, H. Communication accommodation theory: Negotiating personal relationships and social identities across contexts. (Cambridge University Press, 2016).

44. Bourhis, R. Y. & Giles, H. The language of intergroup distinctiveness. Language, ethnicity and intergroup relations 13, 119 (1977).

45. Fulcher, B. D., Little, M. A. & Jones, N. S. Highly comparative time-series analysis: the empirical structure of time series and their methods. Journal of the Royal Society Interface 10, 20130048 (2013).

46. Fulcher, B. D. & Jones, N. S. hctsa: A computational framework for automated time-series phenotyping using massive feature extraction. Cell systems 5, 527–531 (2017).

47. Paul, A., McLendon, H., Rally, V., Sakata, J. T. & Woolley, S. C. Behavioral discrimination and time-series phenotyping of birdsong performance. PLOS Computational Biology 17, e1008820 (2021).

48. Javed, A., Lee, B. S. & Rizzo, D. M. A benchmark study on time series clustering. Machine Learning with Applications 1, 100001 (2020).

49. Takahashi, D. Y. et al. The developmental dynamics of marmoset monkey vocal production. Science 349, 734–738 (2015).

