## Supplementary materials for "Dynamic vocal learning in adult marmoset monkeys"

**The PDF file includes:**

Figs. S1 to S4
Tables S1 and S2
References 1 and 2

**Other Supplementary Materials for this manuscript include the following:**

Movies S1 and S2

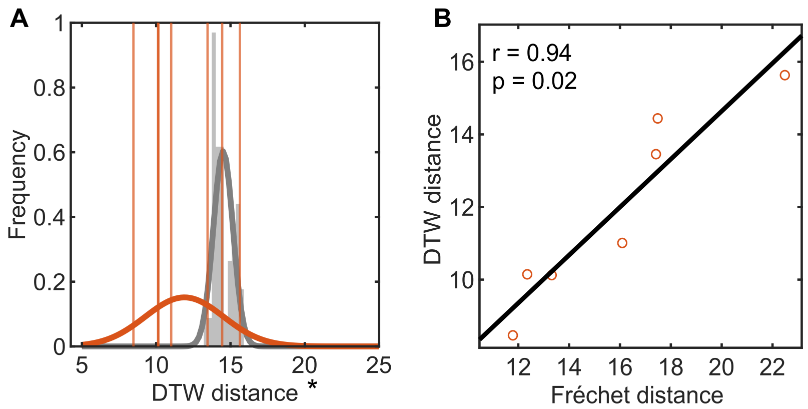


**Fig. S1. Dynamic Time Warping (DTW) distance measures of dyadic acoustic synchrony are highly correlated to Fréchet distance measures.** (**A**) Acoustic feature synchrony in n=7 actual pairs (orange) compared to n=42 control pairings (grey) measured using DTW distances between the trajectories of the marmoset male and the female in a pair. Orange vertical lines depict the DTW distance values of actual pairs. Grey (normalized) histograms depict the distribution of DTW distance values of control pairs. Orange and grey curves are Gaussian probability distribution function fits to actual pairs and random pairs respectively. Lower values indicate better synchrony. **p<0.01, *p<0.05, two-tailed Welch's t-test. (**B**) Pearson's correlation between Fréchet and DTW distances for the n=7 actual pairs.

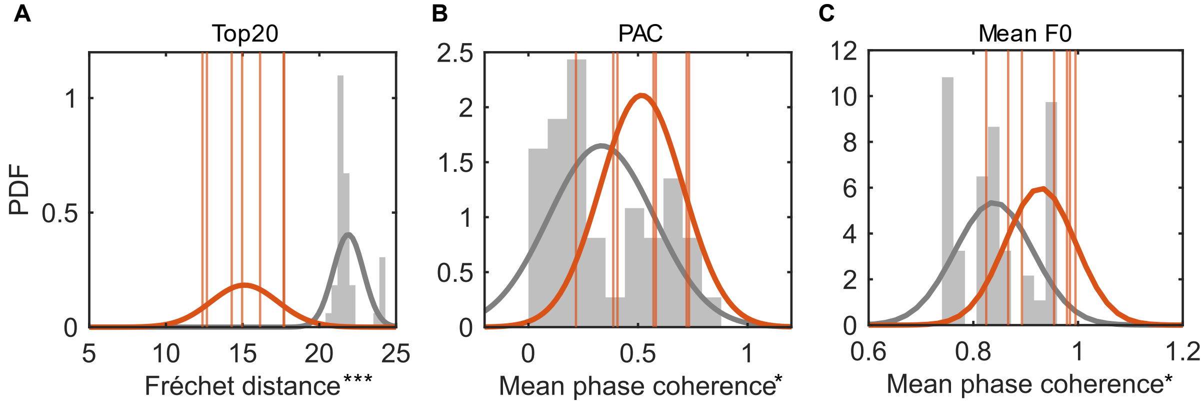


**Fig. S2. Dyadic acoustic feature synchrony is also apparent when convergence is not accounted for or other spectral features are used.** (**A**) Synchrony in the top20 acoustic features in n=7 actual pairs (orange) compared to n=42 control pairings (grey) measured using Fréchet distances between the trajectories of male and female marmosets in a pair without controlling for the convergence in values. Orange vertical lines depict the Fréchet distance values of actual pairs. Grey (normalized) histograms depict the distribution of Fréchet distance values of control pairs. Orange and grey curves are Gaussian probability distribution function fits to actual pairs and random pairs respectively. Lower values indicate better synchrony. ***p<0.001, two-tailed Welch's t-test. (**B, C**) Mean phase coherence of the Principal Acoustic Component (PAC) of mean fundamental frequency, mean spectral entropy, frequency of amplitude modulation and call duration (**B**) , and mean fundamental frequency alone (**C**) of trills of n=7 actual pairs (orange) compared to n=42 control pairings (grey). Orange vertical lines depict the mean phase coherence values of actual pairs. Grey (normalized) histograms depict the distribution of mean phase coherence values of control pairs. Orange and grey curves are Gaussian probability distribution function fits to actual pairs and random pairs respectively. Higher values indicate better synchrony. *p<0.05, two -tailed Welch's t-test.

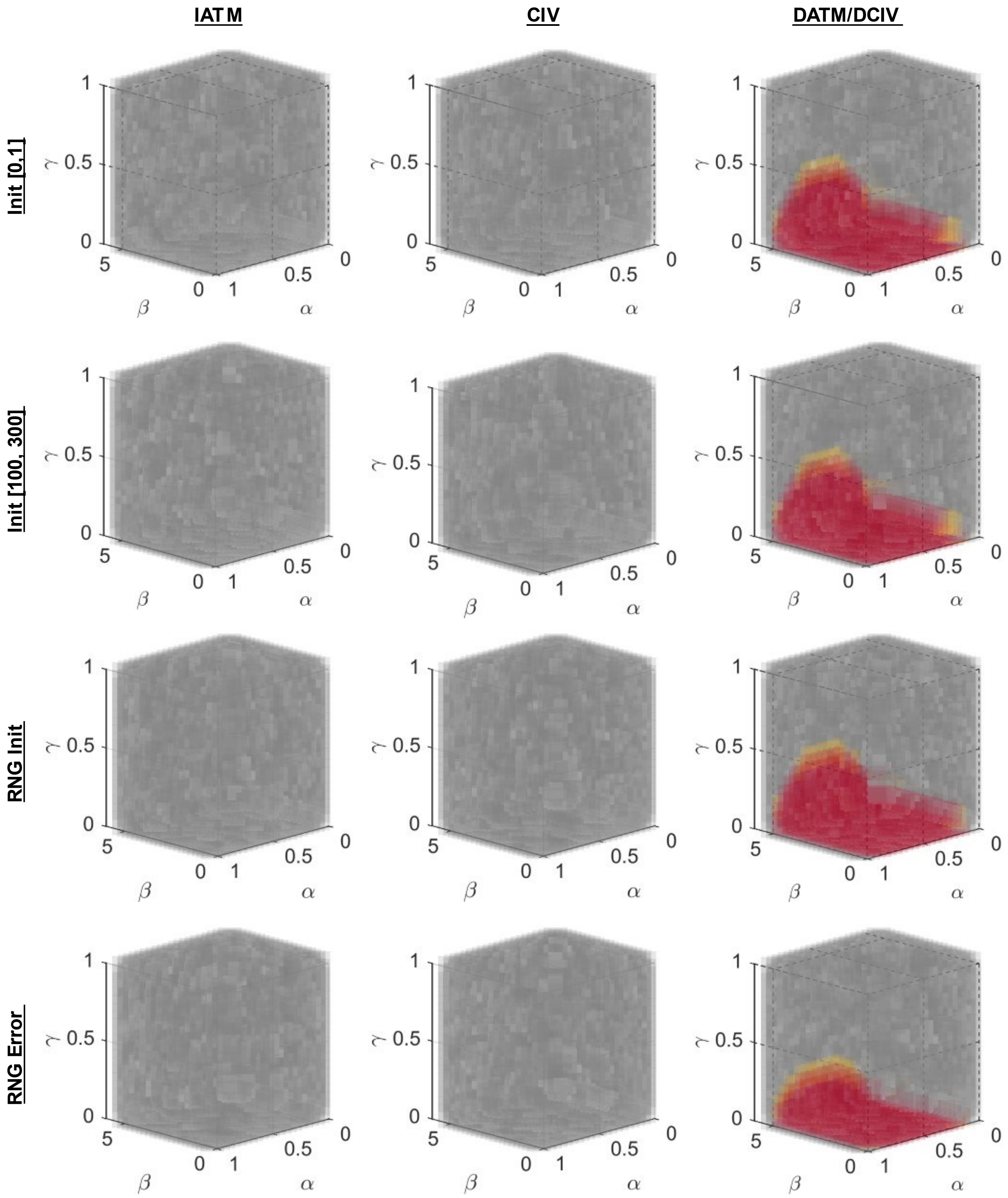


**Fig. S3. Models with *E = f(t)* error term are robust to changes in acoustic feature initialization and random number generators.** Columns correspond to different models (IATM, CIV, DATM/DCIV) and rows correspond to model initialization conditions. Each plot depicts the model parameter space. Model parameter values for which the acoustic features of the n=7 virtual pairs are significantly more synchronized than the n=42 control pairs are marked in yellow if $p<0.05$, and in red if $p<0.01$ (two-tailed Welch's t-test, acoustic trajectory similarity calculated using Fréchet distance). The rest of the points are filled in grey. The fill colors are made slightly transparent such that points behind them are visible. α is the learning rate while β and γ are error coefficients. Abbreviations: IATM = Initial Auditory Template Matching, CIV = Convergence to Intermediate Value, DATM = Dynamic Auditory Template Matching , DCIV = Dynamic Convergence to Intermediate Value, Init [0, 1] = acoustic feature values initialized randomly from a uniform distribution between 0 and 1, Init [100, 300] = acoustic feature values initialized randomly from a uniform distribution between 100 and 300, RNG Init = different random number generator seed for acoustic feature value initialization, RNG Error = different random number generator seed for gaussian noise in the error term.

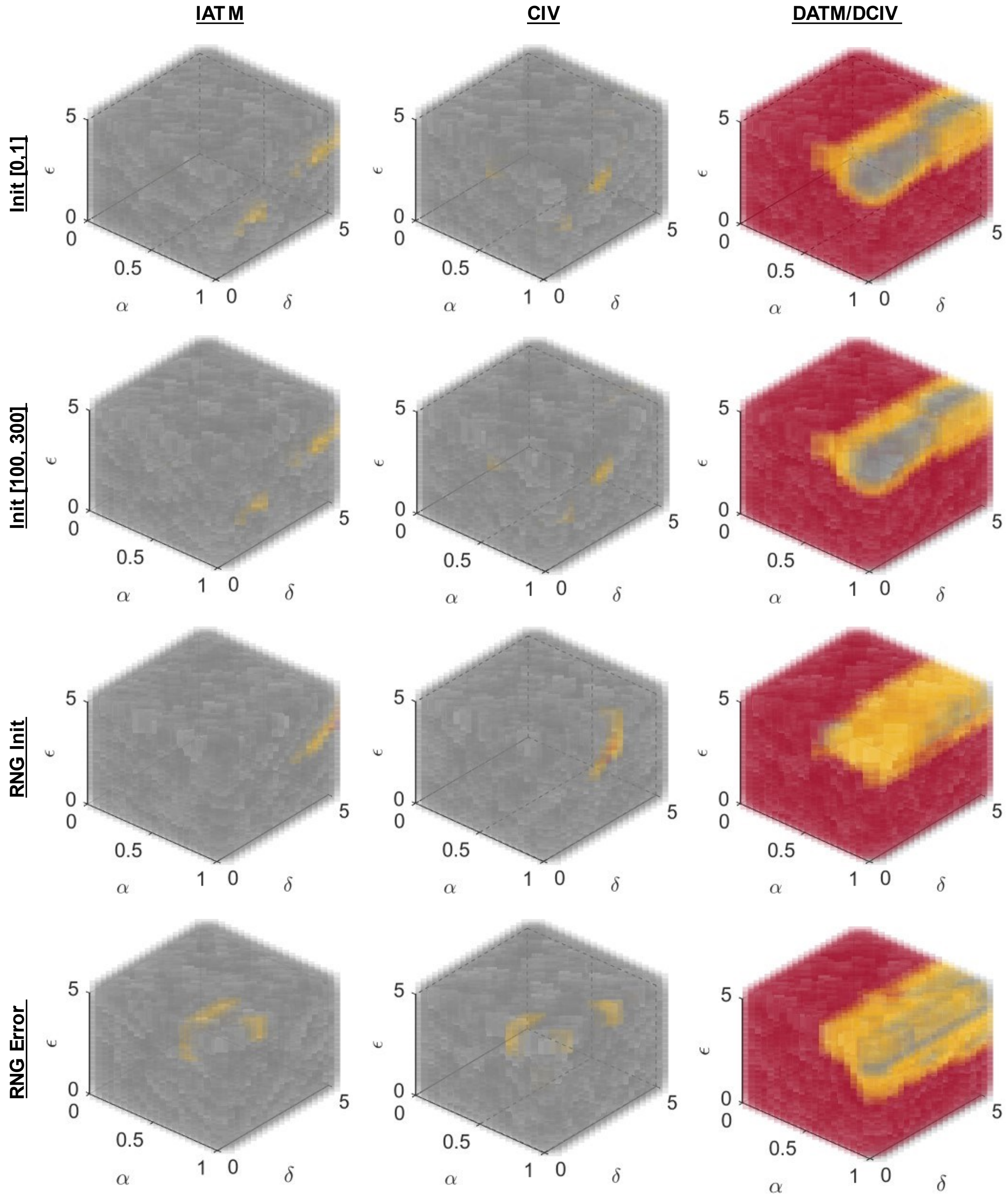


**Fig. S4. Models with *E = f(dv/dt)* error term are robust to changes in acoustic feature initialization and random number generators.** Columns correspond to different models (IATM, CIV, DATM/DCIV) and rows correspond to model initialization conditions. Each plot depicts the model parameter space. Model parameter values for which the acoustic features of the n=7 virtual pairs are significantly more synchronized than the n=42 control pairs are marked in yellow if $p<0.05$, and in red if $p<0.01$ (two-tailed Welch's t-test, acoustic trajectory similarity calculated using Fréchet distance). The rest of the points are filled in grey. The fill colors are made slightly transparent such that points behind them are visible. α is the learning rate while δ and ε are error coefficients. Abbreviations: IATM = Initial Auditory Template Matching, CIV = Convergence to Intermediate Value, DATM = Dynamic Auditory Template Matching , DCIV = Dynamic Convergence to Intermediate Value, Init [0, 1] = acoustic feature values initialized randomly from a uniform distribution between 0 and 1, Init [100, 300] = acoustic feature values initialized randomly from a uniform distribution between 100 and 300, RNG Init = different random number generator seed for acoustic feature value initialization, RNG Error = different random number generator seed for gaussian noise in the error term.

**Table S1. Top 20 features extracted from marmoset trills.** Ranks and descriptions of Highly Comparitive Time Series Analysis (HCTSA) functions used to extract top 20 features from trills. Descriptions taken from Fulcher et al.[1]. Table adapted from Phaniraj et al.[2].

| Feature rank | HCTSA function | Description |
|---|---|---|
| 1 | MF-arfit-1-8-sbc-sumA | Autoregressive model fitting. Optimum order is chosen from the range 1 to 8. Detects inherent repeating patterns. |
| 2 | SY-SlidingWindow-mom3-ent10-10 | Time series cut into 10 equal windows. Skewness calculated for each window. Entropy of skewness values given as output. |
| 3 | SY-RangeEvolve-nuql1000 | Describes how the range of the time series (i.e range of pressure values in the acoustic waveform) changes with time. |
| 4 | EX-MovingThreshold-1-002-stdkickf | Tracks the number of extreme events using a dynamic 'moving threshold'. Every new point in the timeseries is classified as either 'extreme' or 'not extreme' based on the history of extreme events. The standard deviation of this threshold across the timeseries is given as output |
| 5 | EX-MovingThreshold-01-002-mediankickf | Tracks the number of extreme events using a dynamic 'moving threshold'. Every new point in the timeseries is classified as either 'extreme' or 'not extreme' based on the history of extreme events. The median of this threshold across the timeseries is given as output. |
| 6 | MF-arfit-1-8-sbc-A4 | Autoregressive model fitting. Optimum order is chosen from the range 1 to 8. Detects inherent repeating patterns. |
| 7 | SY-SpreadRandomLocal-200-100-stdskew | 100 segments each 200ms long are randomly selected, skewness calculated, and the standard deviation of the skewness values given as output. |
| 8 | SB-MotifThree-diffquant-ccaa | Performs coarse graining of the time series. |
| 9 | SY-LocalGlobal-std-p5 | Compares the standard deviation of the first 1/5th of the time series to that of the entire time series |
| 10 | PH-Walker-momentum-2-sw-taudiff | A hypothetical particle that moves along the length of the time series is simulated. The movement is dynamic and depends on the pressure value of the acoustic waveform at that time. The trajectory of the particle is summarised and compared to that of the actual timeseries. |
| 11 | IN-AutoMutualInfoStats-40-gaussian-pcrossmedian | Median value of the auto-mutual information |
| 12 | SC-MMA-0--n5-5-maxHurstExponent | Multifractal scaling of the time series performed. Maximum Hurst exponent given as output |
| 13 | TSTL-localdensity-5-40-ac-2-minden | Provides estimates of local densities while performing time-delay embedding |
| 14 | NL-TSTL-ReturnTime-10-1-1-n1-ac-8-iqr | Time taken for the pressure value of the acoustic waveform to return to the same location. Provides evidence for periodicities in the data. |
| 15 | SY-SpreadRandomLocal-100-100-stdsampen1-015 | 100 segments each 100ms long are randomly selected, sample entropy calculated, and standard deviation of those values returned. |
| 16 | SC-MMA-0--n5-5-qHurstTrend | Multifractal scaling of the time series performed. $5^{th}$ quantile of Hurst exponent given as output |

| | | |
|---|---|---|
| 17 | CO-TranslateShape-rectangle-2-pts-ones | A rectangle is moved along the length of the time series and statistics performed on all points lying inside the rectangle. |
| 18 | CO-TranslateShape-rectangle-2-pts-fives | A rectangle is moved along the length of the time series and statistics performed on all points lying inside the rectangle. |
| 19 | SC-MMA-0--n5-5-meanHurstExponent | Multifractal scaling of the time series performed. Mean Hurst exponent given as output |
| 20 | SY-SpreadRandomLocal-200-100-stdac2 | 100 segments each 200ms long are randomly selected, autocorrelation coefficient calculated, and the standard deviation of the autocorrelation coefficients values given as output. |

**Table S2. Range of model parameters used for simulations.**

| Parameter | Range | Justification |
|---|---|---|
| Learning rate α | (0, 1] | At α=0, no convergence/learning takes place. Therefore, positive non-zero values were chosen. At α=1, the amount of vocal change in 1 iteration is equal to the initial distance between the vocalizations of the 2 individuals, meaning the individuals switch their vocalizations in 1 iteration. This was set as the upper limit. |
| Error coefficient β | (0, 5] | Entire error term vanishes at β=0. Therefore β>0 was chosen. At β=5 the standard deviation of the Gaussian distribution from which the random error is calculated is 5 times the initial vocal distance between the vocalizations of the 2 individuals which is sufficiently large. |
| Error coefficient γ | (0, 1] | At γ=0, the error term does not decay with time. At γ=1, the error term reduces 1/e times in every iteration. This is a sufficiently steep decay in the error term as dyadic acoustic synchrony vanishes in all models at γ>0.5. |
| Error coefficient δ | (0, 5] | At δ=0, the error term vanishes when no vocal change occurs. At δ=5 a constant error term remains in the form of a random error from a Gaussian distribution with standard deviation 5 times the initial vocal distance between the vocalizations of the 2 individuals, which is sufficiently large. |
| Error coefficient ε | (0, 5] | At ε=0, the error term is a constant. At ε=5, the error term is 5 times the amount of vocal change undergone by the individual in the previous day, which is sufficiently large. |

**Movie S1. An example of 3D visualization of dyadic acoustic feature synchrony in a marmoset pair.** The movie depicts the trajectories of trills of a male (joined by an orange line) and a female (joined by a purple line) in an acoustic feature space comprising of 3 principal components (PCA performed on 20-dimensional acoustic feature space) after pairing. Number of days after pairing is depicted on top of the video. Trajectories were obtained post detrending to dissociate vocal convergence and acoustic feature synchrony.

**Movie S2. An example of 3D visualization of dyadic acoustic feature synchrony in a virtual pair simulated by the model.** The movie depicts the trajectories of trills of 2 virtual individuals in an acoustic feature space comprising of 3 principal components (PCA performed on 20-dimensional simulated acoustic feature space). The iteration number (equivalent to days after pairing) is depicted on top of the video. Trajectories were obtained post detrending to dissociate vocal convergence and acoustic feature synchrony in the model.

**References for supplementary materials**


1. Fulcher, B. D. & Jones, N. S. hctsa: A computational framework for automated time-series phenotyping using massive feature extraction. *Cell systems* **5**, 527–531 (2017).
2. Phaniraj, N., Wierucka, K., Zürcher, Y. & Burkart, J. M. Optimising source identification from marmoset vocalisations with hierarchical machine learning classifiers. 2022.11.19.517179 Preprint at https://doi.org/10.1101/2022.11.19.517179 (2022).
